# A Simple way to Improve a Conventional A/O-MBR for High Simultaneous Carbon and Nutrients Removal from Synthetic Municipal Wastewater

**DOI:** 10.1101/590042

**Authors:** Adoonsook Dome, Chia-Yuan Chang, Wongrueng Aunnop, Pumas Chayakorn

## Abstract

In this study, two anoxic-oxic-MBR systems (conventional and biofilm) were operated in parallel under complete SRT to compare system performance and microbial community composition. Moreover, with the microbial communities, comparisons were made between adhesive stage and suspended stage. High average removal of COD, NH_4_^+^-N and TN was achieved in both systems. However, TP removal efficiency was remarkably higher in BF-A/O-MBR when compared with C-A/O-MBR. TP mass balance analysis suggested that under complete SRT, sponges play a key role in both phosphorus release and accumulation. The qPCR analysis showed that sponge biomass could maintain higher abundance of total bacteria than suspended sludge. Meanwhile, AOB and denitrifiers were enriched in the suspended sludge rather than the sponge biomass. Results of pyrosequencing reveal that the compacted sponge in BF-A/O-MBR could promote the growth of bacteria involved in nutrient removal and reduce the filamentous and bacterial related to membrane fouling in the suspended sludge.

## Introduction

The increase in nutrients, especially nitrogen and phosphorus, in domestic wastewater treatment plant discharge can cause cultural eutrophication in surface waters. The manifestations of these phenomena are known as algal blooms that occur during the warmest season. Excessive nutrient concentration can accelerate the growth of microorganisms (including algae) and other aquatic plants in the receiving waters, leading to low dissolved oxygen concentrations. Due to the fact that conventional biological treatment processes have been designed to meet secondary treatment effluent standards that typically do not remove nitrogen and phosphorus to the extent needed to protect the receiving waters, wastewater treatment systems are required to achieve processes that can enhance the efficiency of biological nutrient removal to safe levels. These processes are referred to as biological nutrient removal (BNR).

Since autotrophic nitrifying bacteria, namely phosphorus accumulating organisms (PAOs), and heterotrophic bacteria compete with each other for habitat and growth under the same operational conditions, the BNR processes normally become unpromising with regard to simultaneously high organic carbon and nutrient removal [1]. Therefore, it has become extremely important to develop reliable technologies that can simultaneously treat organic carbon and nutrients in municipal wastewater. Several research studies have attempted to combine the advantages of attracting growth biofilm and the membrane bioreactor (MBR) process in order to achieve some of the boundaries of traditional MBR that have been based on the activated sludge process. The application of attracting biofilm in MBR could be achieved by the addition of media (e.g., biofilm carriers) in moving or fixed bed configurations, or by the addition of aerated membranes in the bioreactor as a form of support (i.e., substratum) for biofilm growth.

The use of media in the aerobic MBR, called biofilm MBR, could be a better alternative to conventional aerobic MBR which may increase the treatment performance by high biomass concentrations and reduce membrane fouling [2]. Additionally, total nitrogen removal is reported to be higher in the biofilm aerobic MBR systems when compared to conventional aerobic MBR systems according to the authors of several published studies [2,3]. Higher total nitrogen (TN) removal rates have been mostly attributed to simultaneous nitrification/denitrification (SND) that takes place in deeper layers of the biofilm component where anoxic conditions have occurred. The sponge carriers seem to provide good SND conditions since they provide anoxic conditions inside the carrier element. However, phosphorus removal has barely been mentioned by those who have studied aerobic biofilm MBR due to the fact that these systems did not provide an anaerobic zone for PAO in utilizing organic carbon and releasing phosphates.

A combination of anoxic and aerobic MBR compartments with sludge recirculation known as A/O-MBR has been studied frequently for carbon and nitrogen removal in wastewater treatment. Although the A/O-MBR system achieves good simultaneous carbon and nitrogen performance, the system was found to be ineffective for the treatment of phosphorus since at least three separate reactors are required to account for the conditions of differing environments (i.e., anaerobic, anoxic and aerobic), as required by the microorganisms needed in the nitrogen and phosphorus removal process. However, the presence of an anaerobic tank in the biological nutrient removal process requires a larger footprint and is more difficult to operate than an anoxic and aerobic tank. Khan et al. [3] suggested that sponges that act as bio-carriers for attached growth media can be divided into two sub-microenvironments based on dissolved oxygen (DO) gradient values. DO concentration tends to decrease from the surface to the inside of the sponge, providing an aerobic zone at the sponge surface for heterotrophic bacteria (such as PAOs) and nitrifying bacteria. For the interior of the sponge, the anoxic/anaerobic zone is created for the purposes of denitrifying bacteria. Therefore, there are only two tanks (anoxic and aerobic) involved in the process. Notably, phosphorus release and uptake would be possible to achieve if the sponge were placed inside the anoxic compartment to create an anaerobic zone for PAOs. In this regard, phosphates are supposed to accumulate in greater amounts in the sludge biomass, instances of dissolved in the wastewater. Consequently, it would not need further treatment via alternative processes such as ion exchange or chemical precipitation in the permeate stream.

The possibility of operating MBR without sludge withdrawal has been explored by several researchers who have mainly focused on the removal of efficiencies and operational aspects [4,5]. All these authors have indicated the biological applicability of complete sludge retention, while reporting high and stable degradation rates and very limited sludge production. However, due to the fact that phosphorus removal is achieved by the discharge of phosphorus-enriched sludge, phosphorus removal has barely been mentioned in the previous experiments with no sludge withdraw. Consequently, popularizing MBR with complete sludge retention in practice still remains in doubt because of inadequate knowledge on the correlation of microbial communities and nutrient removal.

The objective of this study is to introduce a simple way to achieve high simultaneous nitrogen and phosphorus removal in the A/O-MBR system by combining biofilm as active biomass that is packaged inside the anoxic compartment. The microbial community structures and compositions between the biofilm anoxic/oxic membrane bioreactor system (BF-A/O-MBR) and the conventional anoxic/oxic membrane bioreactor system (C-A/O-MBR), along with a potential link to the microbial community change to the nutrient biodegradation, were investigated to clearly understand why biofilm that is coupled with A/O-MBR is better than the conventional A/O-MBR system in terms of the biological removal by microorganisms. Moreover, within the microbial community, the difference between the adhesive stage and the suspended stage in the BF-A/O-MBR was also investigated. The microbial community that is present in the sponge biomass of the BF-A/O-MBR system and the suspended sludge of both systems was analyzed by pyrosequencing. An effort has been made to compare treatment efficiency levels and the microbial community compositions of both systems under complete sludge retention.

## Materials and Methods

### Lab-scale anoxic and aerobic membrane bioreactor

The study was carried out using two lab-scale anoxic and aerobic membrane bioreactors (BF-A/O-MBR and C-A/O-MBR). The reactors were working in parallel and fed with synthetic domestic wastewater (Fig 1). Each of the two A/O-MBR systems used in this experiment consisted of two reactors with a total operating volume of 4.5 l (1.5 l for the anoxic tank and 3.0 l for the aerobic MBR tank). In the aerobic MBR tank of each system, a polyvinylidene fluoride (PVDF) hollow-fiber membrane module with a pore size of 0.1 µm and a total membrane area of 0.025 m^2^ was installed. Air diffusers were constructed beneath the membrane modules to continuously supply oxygen for biomass growth and to constantly scour the membrane surface for potential fouling control. The dissolved oxygen (DO) concentrations in the anoxic tank and the aerobic MBR tank were controlled at approximately 0.3 mg/l and 2.0 mg/L, respectively. A pressure gauge was installed to measure the transmembrane pressure (TMP) during the filtration period.

**Fig 1.**
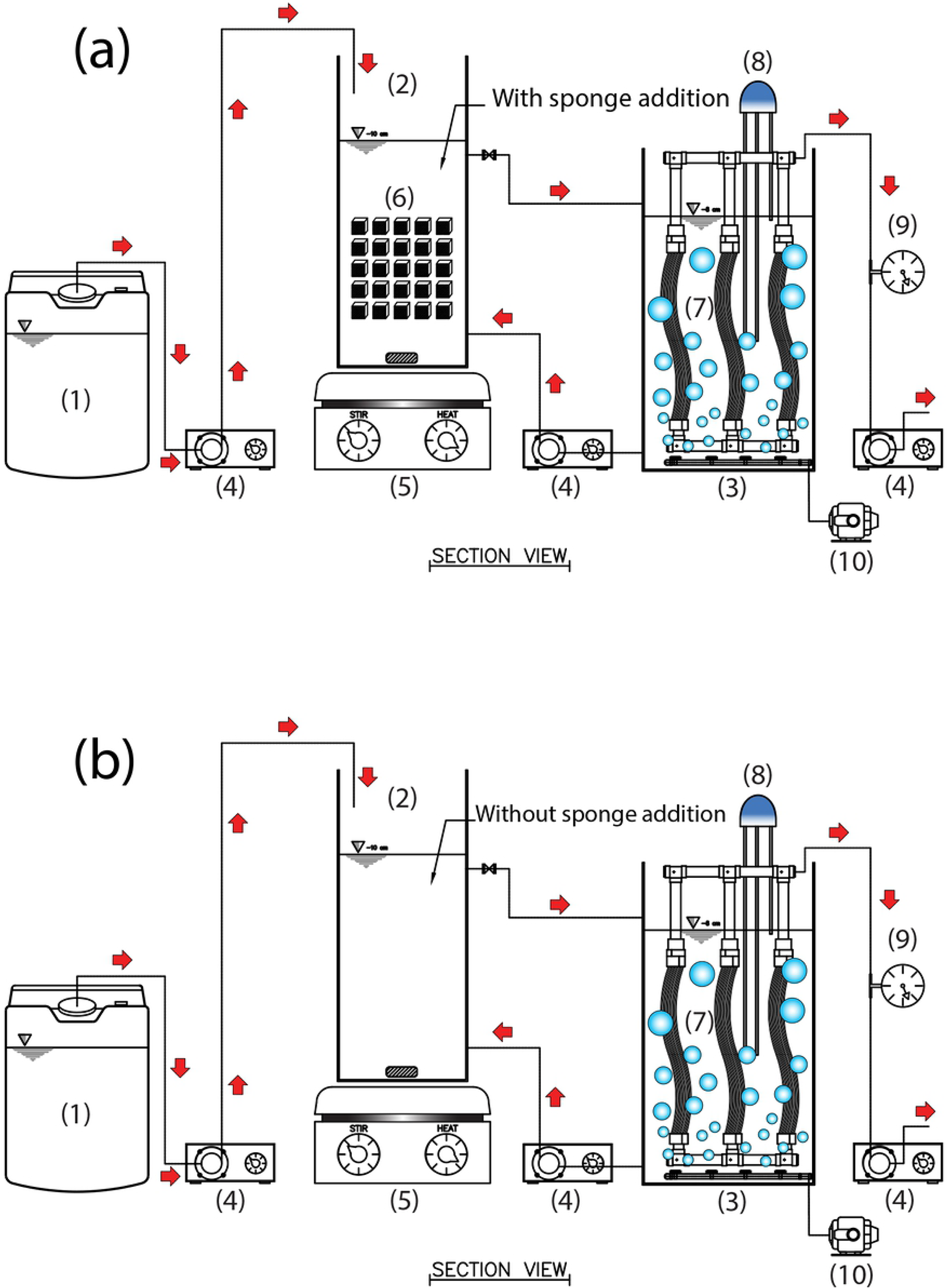
The schematic diagram of the laboratory-scale A/O-MBR system. System (a) (with compacted sponge addition) and system (b) (without compacted sponge addition) consisted: (1) feed tank; (2) anoxic tank; (3) MBR tank; (4) pump; (5) stirrer; (6) sponges; (7) membrane module; (8) level sensor; (9) Pressure gauge; (10) air pump.

As for the BF-A/O-MBR system, 50 pieces of one-cubic-centimeter-sponge were compacted inside the anoxic tank with a 10% volume fraction of the anoxic tank. In both systems, synthetic wastewater from the feed tank was introduced to the anoxic tank using a peristaltic pump (Model no. 7553-71, Masterflex). The anoxic effluent was made to flow to the aerobic MBR tank using gravity. A water level controller was installed in the aerobic MBR tank of each system to maintain a constant water level corresponding to a total hydraulic retention time (HRT) of 9 h (3 h for anoxic tank and 6 h for aerobic MBR tank). A permeate flux of 20 L/m^2^.h with a permeate flow rate of 14.4 l/day was maintained during the entire experimental procedure. Intermittent permeation with 5 min suction and 1 min pause was applied to minimize membrane fouling. The system was operated at ambient temperatures. The suspended sludge in the aerobic MBR tank was recirculated to the anoxic tank by a return pump with a recirculation rate of 2 times of the influent flow rate. It is important to note that all of the systems were operated without sludge withdrawal except when it was necessary to analyze certain relevant parameters.

The raw wastewater employed in this study was synthetic domestic wastewater imitated from domestic wastewater characteristics in Taiwan with concentrations of 210 ± 5 mg/L chemical oxygen demand (COD), 60 ± 1 mg/L total nitrogen (TN), and 5.5 mg/L total phosphorus (TP). The components of the synthetic wastewater were as follows: C_2_H_5_COONa (200 mg/l), NH_4_Cl (200 mg/l), KH_2_PO_4_ (28 mg/l), NaHCO_3_ (220 mg/l), MnCl_2_.4H_2_O (0.19 mg/l), ZnCl_2_.2H_2_O (0.0018 mg/l), CuCl_2_.2H_2_O (0.022 mg/l), MgSO_4_.7H_2_O (5.6 mg/l), FeCl_3_.6H_2_O (0.88 mg/l), and CaCl_2_.2H_2_O (1.3 mg/l). The reactors were inoculated with activated sludge from a domestic wastewater treatment plant located at An-Ping Tainan, Taiwan. A sample of the activated sludge collected from the aeration basin was settled for 4 h, and the supernatant was discarded. At the start of each system, a portion of activated sludge was diluted with DI water to 10 l, aerated in the container and fed with a stock solution at an influent flow rate of 3.5 l/day to restore its activity. After 7 days, a 4.5 l volume of the renewed sludge was applied as an inoculum to maintain an initial MLSS concentration of approximately 5,000 mg/l.

### Analytical method for wastewater quality parameters

Mixed liquor suspended solids (MLSSs), mixed liquor volatile suspended solids (MLVSSs), chemical oxygen demand (COD), ammonia nitrogen (NH_4_^+^-N), nitrite (NO_2_ ^-^-N), nitrate (NO_3_ ^-^-N), total nitrogen (TN), and total phosphorus (TP) concentrations were measured according to the standard methods [6]. Dissolved oxygen and pH were determined using a WTW Multi pH/Oxi 340i dissolved oxygen meter. A differential pressure gauge was used to measure the transmembrane pressure (TMP) of the membrane module.

Samples of mixed liquor for EPS analysis were taken from the anoxic tank and the aerobic MBR tank. All samples were cooled to 4°C to mitigate microbial activity. The samples were then centrifuged at 6,000 g for 20 min at a temperature of 4°C. The supernatant was filtrated with a 0.45 µm membrane filter. The precipitation was re-suspended in the buffer solution with the same volume of the supernatant. The extraction of EPS was based on the cation exchange resin (Dowex^®^ Marathon^®^ C, Na^+^ form, Sigma-Aldrich, Bellefonte, PA) extraction method, as has been described by Frølund et al. [7]. The exchange resin of 70 g of Dowez/g MLVSS was added to 30 ml of the sample. The sample and the exchange resin were then mixed at 600 rpm (4°C) using a magnetic stirrer for 2 h. After that, the samples were centrifuged for 15 min at 12,000 g in order to remove the suspended solids. The supernatant was then filtered through a 0.22 µm membrane filter. The filtrate was presented as the total EPS. The carbohydrate content and the protein content were measured using the phenol-sulfuric acid method and the Modified BSA kit based on the methods described by Dubois et al. [8] and Lowry et al. [9], respectively. Glucose and bovine serum albumin (BSA) were used as the carbohydrate standard and the protein standard, respectively. The sum of the carbohydrate and the protein EPS concentration was presented as the total EPS concentration.

### Sampling and DNA extraction

The sponge biomass of the BF-A/O-MBR system and the sludge biomass in the aerobic MBR tank of both systems were sampled periodically throughout a 45-day period. DNA extraction was performed using the Fast-DNA SPIN kit for soil (MP Biomedicals, Solon, OH, USA) with some modifications of the protocol. This method used a mix of chemicals and physical methods. The lysis process is the initial process that involves adding lysis chemical into the sample inside a tube that contains beads and is then put throgh a physical high-speed mixer. After the lysis procedure, the experimental procedure continues with several additional steps. The removal of humic acid, cell debris, and protein was achieved through the use of chemicals and a filtration step. In the last purification procedure, all washing chemicals were removed. The final step involves eluting DNA into a clean tube until it was ready to be used. The extracted DNA was evaluated on 1% (wt/vol) agarose gel and stored at −20°C until further use.

### Quantitative real-time PCR assays

For each sample, a quantitative real-time PCR was carried out in duplicate with a LC 480 SYBR Green QPCR Master Mix (Roche Diagnostics, Mannheim, Germany) in a light Cycler 2.0 system instrument. The quantification of bacterial AOB *amoA* genes, denitrifying *Nirs* genes, and total bacterial 16S rRNA genes were performed using the primers (Supplementary Table 1S, supporting information). The 10-fold dilution series of standard calibration ranged from 5.0 to 5×10^7^ copies/reaction. The PCR mixture at a volume of 20 µl contained the reagent as is shown in Table 2S (supporting information). The DNA template was changed to ddH_2_O when conducting the negative control. The PCR mixture was the same for all genes.

**Table 1.** Influent concentration, effluent concentration and system performance of BF-A/O-MBR system and C-A/O-MBR system.

| System types | BF-A/O-MBR | C-A/O-MBR |
| --- | --- | --- |
| Influent COD ($\text{mg}\cdot\text{l}^{-1}$) | $246.65 \pm 9.0$ | $242.06 \pm 8.6$ |
| Effluent COD ($\text{mg}\cdot\text{l}^{-1}$) | $7.44 \pm 2.9$ | $4.07 \pm 2.0$ |
| COD removal efficiency (%) | $96.98 \pm 1.2$ | $98.32 \pm 0.8$ |
| Influent $\text{NH}_4^+$ ($\text{mg}\cdot\text{l}^{-1}$) | $59.93 \pm 7.4$ | $56.76 \pm 4.4$ |
| Effluent $\text{NH}_4^+$ ($\text{mg}\cdot\text{l}^{-1}$) | $0.38 \pm 0.1$ | $0.57 \pm 0.02$ |
| $\text{NH}_4^+$ removal efficiency (%) | $99.39 \pm 0.2$ | $98.92 \pm 0.1$ |
| Influent TN ($\text{mg}\cdot\text{l}^{-1}$) | $60.14 \pm 7.4$ | $58.38 \pm 5.0$ |
| Effluent TN ($\text{mg}\cdot\text{l}^{-1}$) | $7.34 \pm 0.7$ | $10.81 \pm 1.0$ |
| TN removal efficiency (%) | $87.79 \pm 2.0$ | $81.48 \pm 2.6$ |
| Influent TP ($\text{mg}\cdot\text{l}^{-1}$) | $5.05 \pm 0.2$ | $5.12 \pm 0.2$ |
| Effluent TP ($\text{mg}\cdot\text{l}^{-1}$) | $0.24 \pm 0.6$ | $3.89 \pm 0.2$ |
| TP removal efficiency (%) | $95.30 \pm 1.2$ | $24.02 \pm 5.3$ |
| MLSS ($\text{mg/l}$) | $13142 \pm 119$ | $9032 \pm 69$ |
| MLVSS ($\text{mg/l}$) | $10910 \pm 201$ | $8530 \pm 136$ |
| EPS protein ($\text{mg/gMLVSS}$) | $77.03 \pm 14.6$ | $41.32 \pm 5.6$ |
| EPS carbohydrate ($\text{mg/gMLVSS}$) | $51.10 \pm 26.4$ | $67.45 \pm 13.8$ |
| Total EPS ($\text{mg/gMLVSS}$) | $128.13 \pm 9.1$ | $108.77 \pm 5.0$ |

**Table 2.** Richness and diversity index of bacteria community in the sponge biomass sample of BF-A/O-MBR system (BF-A/O-MBR (SP)) and suspended sludge samples of both BF-A/O-MBR and C-A/O-MBR system (BF-A/O-MBR (SS) and C-A/O-MBR (SS), respectively).

| Sample ID | BF-A/O-MBR (SP) | BF-A/O-MBR (SS) | C-A/O-MBR (SS) |
| --- | --- | --- | --- |
| Observed OUT | 671 | 623 | 579 |
| Shannon index (H') | 4.78 | 4.36 | 3.95 |
| Species richness (Hmax) | 6.50 | 6.43 | 6.36 |
| Species evenness (E) | 0.69 | 0.68 | 0.62 |
| 1/Simpson index | 37.72 | 36.98 | 24.93 |

### Bacterial community analysis using 454 pyrosequencing

The sponge biomass samples of the BF-A/O-MBR system and the sludge biomass samples of both systems on day 45 were chosen for pyrosequencing analysis. PCR amplification was carried out immediately after DNA extraction, using primers targeting the V3 and V4 regions of the 16S rRNA gene with the forward primer 341F (5’-CCTACGGGAGGCAGCAG-3’) and the reverse primer 805R (5’-GACTACCAGGGTATCTAATCC-3’), which used the extracted DNA as a template. The PCR mixture was composed of 0.8 µl for each forward and reverse primer (10µM, Metabion, Germany), 3 µl of the template DNA for the samples, and 12.5 µl of 1x of Hot Master Mix (Promega GoTaq® Green Master Mix) to a final volume of 25 µl. For the negative control, 3 µl of the elution solution was used. The amplifications were performed under the following conditions: initial denaturation at 95°C for 2 minutes, followed by 30 cycles of denaturation at 95°C for 30 seconds, primer annealing at 60°C for 30 seconds, and extension at 72°C for 30 seconds, with a final elongation at 72°C for 5 minutes.

Sequencing was carried out by using paired-end Illumina MiSeq sequencing on an Illumina MiSeq device (Illumina Inc., San Diego, CA, USA) with 600 cycles (300 cycles for each paired read and 12 cycles for the barcode sequence) according to the manufacturer’s instructions. To artificially increase genetic diversity, it has become common practice to mix in a control library of genomic DNA from the phage phix to prevent focusing and phasing problems due to the sequencing of “low diversity” libraries. Sequence analysis was conducted using the 16S-based metagenomics workflow of MiSeq Reporter version 2.6.2.3 (Illumina). Samples were gathered into a single library for sequencing on Illumina MiSeq sequencing system which generated paired 300 bp reads. Sequences were then demultiplexed based on index sequences. FASTQ files with Quality Score Encoding were created. OTUs clustering and classification at several taxonomic levels, kingdom, phylum, class, order, family, genus, and species, were performed.

### Sequence analysis

The diversity of the microbial community was quantified by Shannon Weiner’s Diversity Index [10]. The ensuing formula was used to calculate the values as:

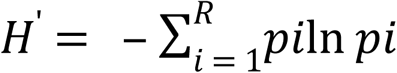

where H′ = Shannon Weiner’s Diversity Index, pi = ratio of individuals in the i species, R = number of species

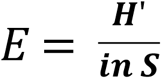

where E = evenness, S = total number of species in the population.

The species richness (H_max_) was calculated as:

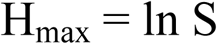

Simpson’s index (D) of the samples was calculated as:

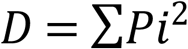

Bacteria assemblage patterns were examined utilizing several approaches including R Project for Statistical Computing version 3.4.2 supported by CRAN (R Core Team, 2017). Sampling runs were grouped based on microbial communities using Hierarchical Cluster Analyses (HCA) and carried out with h cluster function of R using Euclidean distance. The complete option was embraced for the clustering as it includes a greater proportion of the information than 2%. The Pearson’s product momentum correlation coefficient was used to estimate the linear correlation between the abundance of the top 37 most abundant OTUs and the two most important environmental variables (nitrogen and phosphorus). Pearson’s coefficient (r_p_) was always between −1 and +1, where −1 means a perfectly negative correlation and +1 indicates a perfect positive correlation, while 0 indicates the absence of a relationship.

## Results and Discussion

### MBRs performance

General performance of the two MBR systems is summarized in Table 1. The COD and NH_4_^-^- N removal performances were higher than 97% and 99%, respectively, when the influent COD and NH_4_^-^-N loads were 0.65 kg-COD/m^3^.d and 0.16 kg- NH_4_^-^-N /m^3^.d, respectively. Since autotrophic nitrifying bacteria are well-known as slow-growing bacteria [11], the extended mean cell residence time (MCRT) would be able to provide nitrifying bacteria that is sufficient enough to ensure effective nitrification. Therefore, the complete SRT applied in this study was very beneficial for the nitrifying bacteria. However, the TN removal efficiency levels were found to be significantly higher (α = 0.05) in the BF-A/O-MBR system (87.79%) than in the C-A/O-MBR system (81.48%). The results indicate that the reactor configuration (i.e., compacted sponge addition) played a role in denitrification and TN removal. The higher level of TN removal efficiency associated with the BF-A/O-MBR system can mostly be attributed to the multifunctional microbial reactions that occurred in the developing sponge biofilm [2], especially with regard to denitrifying bacteria and DNPAOs, which play crucial roles in nitrogen removal. This aspect will be proven using real-time PCR and pyrosequencing analysis in the next sections.

Surprisingly, the extremely high phosphorus removal performance was observed in the BF-A/O-MBR system with an average removal efficiency of 95.30%, while the average phosphorus removal performance was lower than 25% in the C-A/O-MBR system. Due to the fact that phosphorus removal is achieved by the discharge of phosphorus-enriched sludge, phosphorus removal has rarely been mentioned in the previous experiments that revealed no sludge discharge. Because both systems operated without sludge withdrawal (except in the analysis), the very high phosphorus removal performance in the BF-A/O-MBR system was unexpected. The TP mass balance showed that the average TP feeding rates were 0.061 and 0.062 g-P/day, and that the average TP releasing rates of around 1.31 and 0.03 g-P/day occurred in an anoxic tank of the BF-A/O-MBR and C-A/O-MBR systems, respectively (Supplementary Table 3S, supporting information). The uptake rates in the aerobic MBR tank in the BF-A/O-MBR and C-A/O-MBR systems were found to be 2.41 and 0.18 g-P/day, respectively. The results indicated that when the sponge biomass was presented in the anoxic tank, it could create anaerobic conditions inside the sponge biomass for the release of phosphates by PAOs. Khan et al. [3] suggested that sponges as bio-carrier attached growth media could be divided into two sub-micro-environments based on the dissolved oxygen (DO) gradient. DO concentration tends to decrease from the surface to the inside of the sponge, providing an aerobic zone at the sponge’s surface for heterotrophic bacteria (such as PAOs) and nitrifying bacteria. For the interior of the sponge, the anoxic/anaerobic zone was created for the purposes of denitrifying bacteria and PAOs.

The mass balance result also showed that most of the TP in the influent was adsorbed somehow into suspended biomass 0.015 g-P/day and sponge biomass 0.044 g-P/day in the BF-A/O-MBR system. Meanwhile, in the C-A/O-MBR system, 0.010 g-P/day was only contained by the suspended biomass which produced 0.048 g-P/day in the permeate stream. Therefore, based on the phosphorus feeding rate, 27% and 72% were accumulated in the suspended sludge and sponge biomass in the BF-A/O-MBR system, respectively. According to researchers, extracellular polymeric substances (EPS) play a crucial role regarding the bio-absorption of pollutants [12,13].

Cloete and Oosthuizen [14] have reported that phosphorus removal in the activated sludge might not only be due to PAO activity but also occurs as a consequence of EPS acting as a phosphorus reservoir. Regarding MLVSS and total EPS concentration, it was found that the value of MLVSS and total EPS concentration were significantly higher in the BF-A/O-MBR system than in the C-A/O-MBR system (Table 1). Yigit et al. [15] reported that higher MLSS levels typically increase the concentration of EPS. Therefore, EPS might be one of the most important factors in the phosphate adsorption by way of suspended biomass and sponge biomass in the BF-A/O-MBR system. However, it remains unclear and information is still lacking that would explain what was truly happening in this phenomenon. The phenomenon would need to be further investigated in future work.

### Real-time quantitative PCR

Quantitative real-time PCR of 16S rRNA genes, bacterial *amoA* genes, and *Nirs* genes was applied to estimate the abundance of total bacteria, ammonium oxidizing bacteria (AOB) and denitrifying bacteria, respectively, in the sponge biomass samples (BF-A/O-MBR system) and suspended sludge samples (BF-A/O-MBR and C-A/O-MBR system) (Fig 2). At the beginning of the experiment, the amounts of total bacteria, AOB and denitrifying bacteria of the suspended sludge samples in the BF-A/O-MBR system were started at approximately 9.3 ×10^2^, 1.4 ×10^1^ and 5.2 ×10^1^ copies/ml of sludge biomass, respectively. Meanwhile, the virgin sponge started without any biomass accumulation. The amounts of total bacteria, AOB and denitrifying bacteria in both sponge biomass and suspended biomass samples gradually increased and fluctuated during the first 16 days of operation. A stabilization in abundance of total bacteria, AOB and denitrifying bacteria in both sponge biomass and suspended biomass samples were observed after day 20 of the operation. At the end of the experiment (day 45), total bacteria, AOB and denitrifying bacteria concentration of the BF-A/O-MBR system were 1.1 ×10^11^, 1.6 ×10^3^ and 4.5 ×10^5^ copies/ml of sponge biomass and 6.0 ×10^8^, 9.9 ×10^5^ and 5.1 ×10^6^ copies/ml of sludge biomass, respectively. During the stabilization stage, the amounts of total bacteria in the sponge biomass were higher than that of the suspended sludge. However, for AOB and denitrifying bacteria, the concentrations were always lower in the sponge biomass when compared with the suspended sludge. The results clearly revealed that a compacted sponge at an adhesive stage could contain higher active biomass than a suspended stage. Nevertheless, AOB and denitrifying bacteria likely preferred to grow under the suspended stage rather than under the adhesive stage. Due to the fact that the compacted sponge was added into the anoxic tank, thus, the inner side of the compacted sponge may lack the oxygen and nitrate source which was used as an electron acceptor for AOB and denitrifying bacteria, respectively, resulting in lower amounts of these bacteria groups in the sponge biomass. Therefore, of the majority of the nitrification and denitrification processes in this study tended to occur in the suspended sludge rather than the sponge biomass.

**Fig 2.**
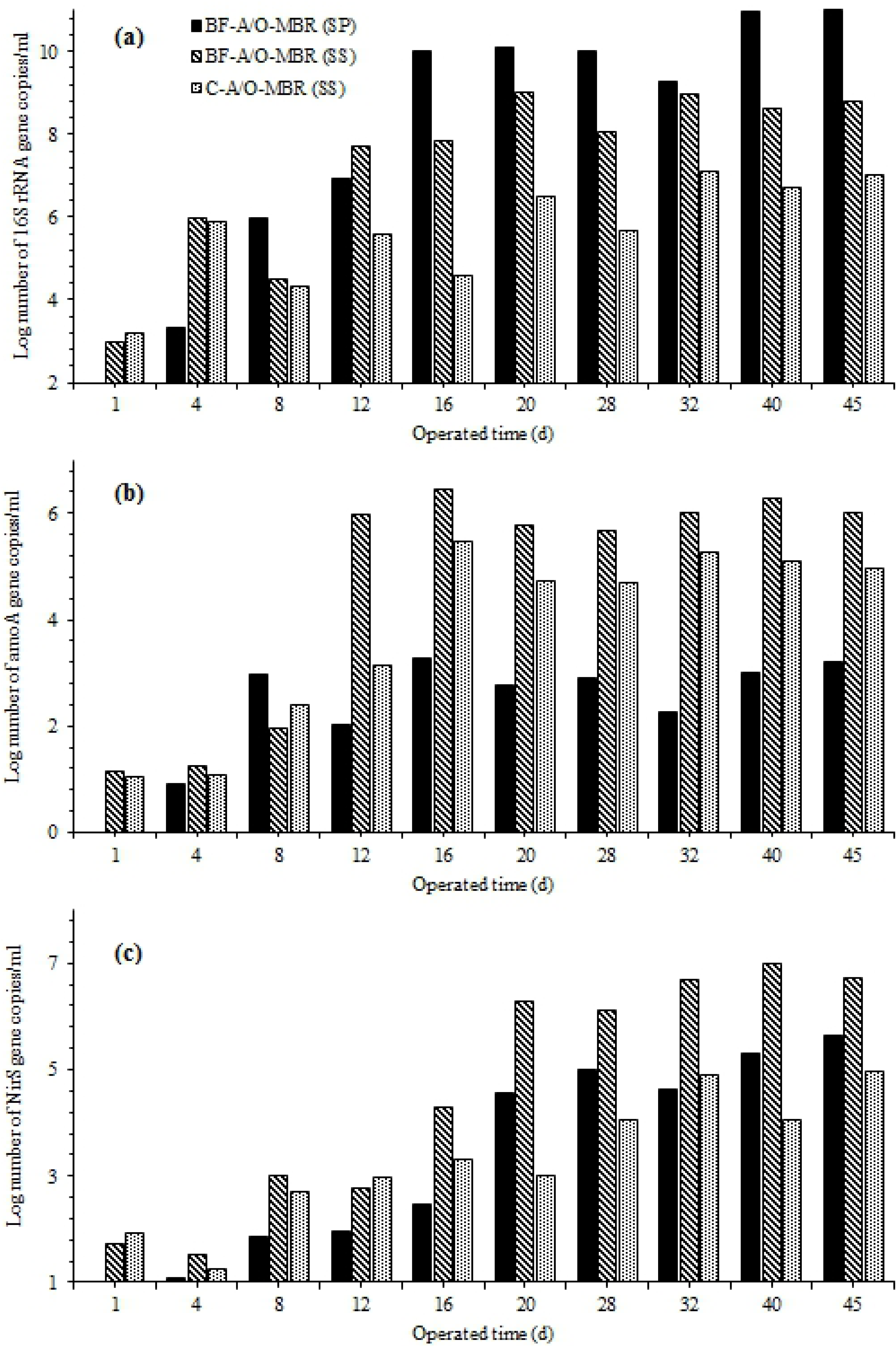
Bacterial 16S rRNA gene (a), AmoA gene (b) and NirS gene (c) copy number change with operating time of sponge biomass sample of BF-A/O-MBR system (BF-A/O-MBR (SP)) and suspended sludge samples of both BF-A/O-MBR and C-A/O-MBR system (BF-A/O-MBR (SS) and C-A/O-MBR (SS), respectively).

When comparing the abundance of the bacteria groups of the two systems (BF-A/O-MBR and C-A/O-MBR systems), it was found that after inoculation, the averages of total bacteria, AOB and denitrifying bacteria of the suspended sludge samples were approximately 1.7 ×10^3^, 1.3 ×10^1^ and 7.0 ×10^1^ copies/ml of the sludge biomass, respectively, in the two reactors. There was no significant difference between the two reactors for the first 8 days of operation (α = 0.05). After day 16, the amounts of total bacteria, AOB and denitrifying bacteria were all higher in the BF-A/O-MBR system. The amounts of total bacteria, AOB and denitrifying bacteria distinctly increased during day 20 in both systems and seemed to be constant after that time. The results suggested that the specific amounts of microorganisms that were chosen for their capability to degrade the chemical component of synthetic wastewater would significantly achieve the stability and performance of both systems after 20 days. At day 45, the concentrations of total bacteria, AOB and denitrifying bacteria were 6.4 ×10^8^, 1.8 ×10^6^ and 5.1 ×10^6^ copies/ml of sludge biomass for the BF-A/O-MBR system and 1.2 ×10^7^, 1.2 ×10^5^ and 9.1 ×10^4^ copies/ml of sludge biomass for the C-A/O-MBR system, respectively. Obviously, the sponge biomass played an essential role in the growth of bacteria in the suspended sludge. Therefore, it could be concluded that the significant improvement of TN removal in the BF-A/O-MBR system, when compared with the C-A/O-MBR system, was achieved mainly by adding a sponge to the anoxic tank.

### Richness and diversity of bacteria community

Diversity indices serve as a valuable tool to quantify diversity in a population and describe its numerical structure, combining richness and evenness components. Two diversity indices, the Shannon index (H’) and the reciprocal of Simpson’s index (D), were used to assess the bacterial community diversity in this study. Simpson’s index is heavily weighted towards the most abundant species in the sample while being less sensitive to species richness. The Shannon index is positively correlated with species richness and evenness and gives more weight per individual than the common species, being sensitive to sample size [16]. Higher numbers indicate greater levels of diversity.

The diversity index results of the sponge biomass sample in the BF-A/O-MBR system and suspended sludge samples in the BF-A/O-MBR and C-A/O-MBR systems are shown in Table 2. Both the Shannon and 1/Simson indexes of 4.78 and 37.72 in the sponge biomass sample showed higher diversity values than those of the suspended sludge sample of the BF-A/O-MBR system at 4.36 and 36.98, respectively. Moreover, the Shannon and 1/Simson indexes of both the sponge biomass and suspended sludge samples in the BF-A/O-MBR system were significantly higher than the C-A/O-MBR system (3.95 and 24.93, respectively). The results showed that the sponge biomass as an active attached biomass was more diverse than the suspended sludge. This finding also revealed that the existence of the compacted sponge in the anoxic tank had an effect on the diversity of the microbial community in the suspended sludge. The reason for this is that when the compacted sponge was constructed in the anoxic tank, the biofilm which developed on the sponge surface became thick and eventually sloughed off to the suspended sludge. Therefore, some of the microorganisms that attached and grew on the sponge surface were exposed to the suspended sludge, which led to a high level of diversity in the suspended sludge.

A Venn diagram of the exclusive and shared OUT (genus level) values of the sponge biomass of the BF-A/O-MBR system and suspended sludge samples of both the BF-A/O-MBR and C-A/O-MBR systems is presented in Fig 3. There were 535 OTUs seen in three samples while 275, 109 and 91 OTUs were unique for the sponge biomass and suspended sludge samples of the BF-A/O-MBR system and the suspended sludge sample of the C-A/O-MBR system, respectively. The difference of unique OTUs in the three samples indicated that the communities in the sponge biomass and the suspended sludge of the system with compacted sponge samples (BF-A/O-MBR system) were more diverse than those in the suspended sludge of the system without a compacted sponge (C-A/O-MBR system).

**Fig 3.**
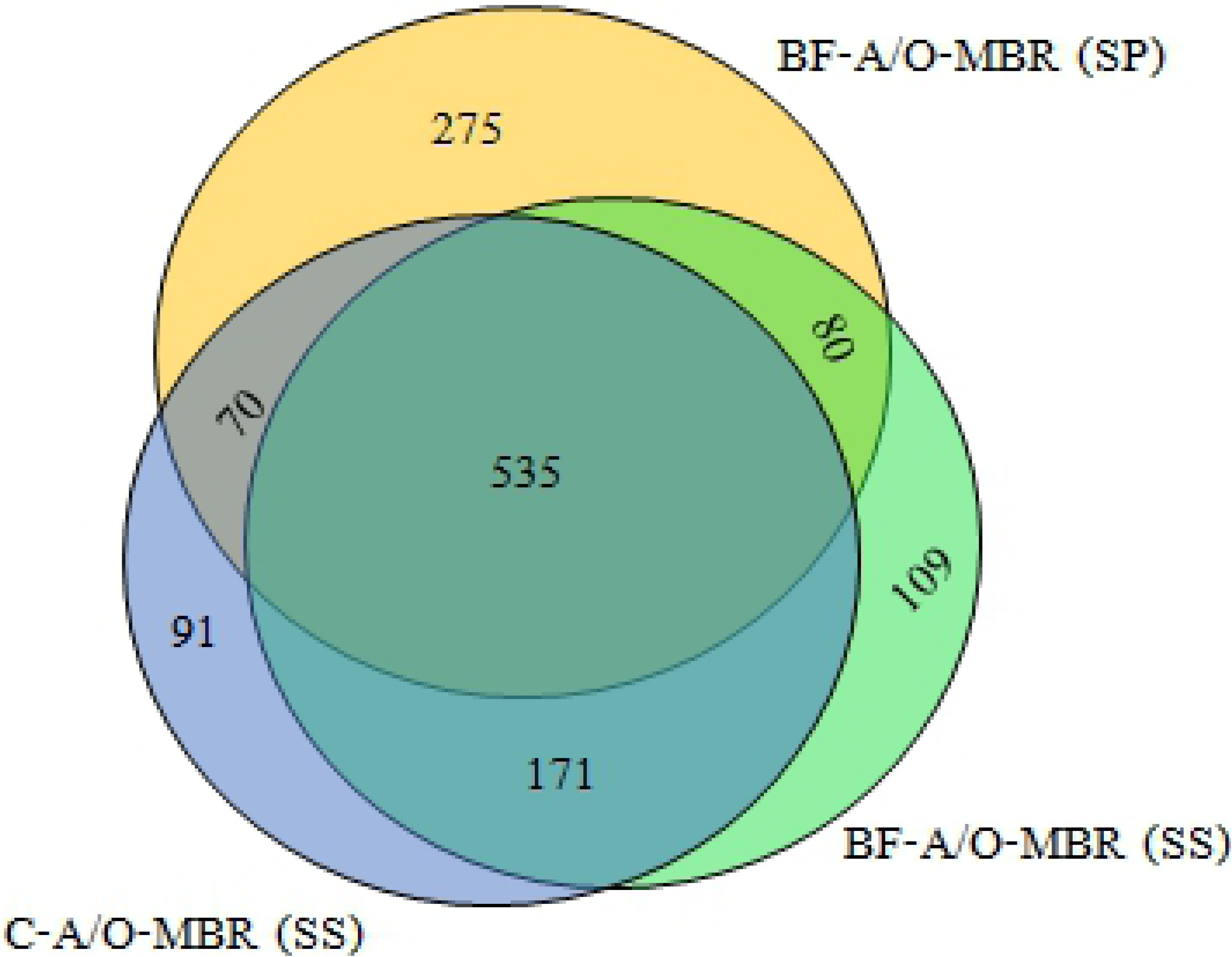
Venn’s diagram of exclusive and shared OUT between sponge biomass sample of BF-A/O-MBR system (BF-A/O-MBR (SP)) and suspended sludge samples of both BF-A/O-MBR and C-A/O-MBR system (BF-A/O-MBR (SS) and C-A/O-MBR (SS), respectively).

Therefore, it could be concluded that the presence of a compacted sponge in the anoxic tank of the BF-A/O-MBR system may contribute to high levels of nitrogen and phosphorus removal performance (87.79% and 95.3%, respectively), which is reflected in the higher presence of microbial diversity in the system. Some of the species (such as nitrifiers, denitrifiers and PAOs) that appeared in the BF-A/O-MBR system and disappeared in the C-A/O-MBR system might be the main microorganisms that are involved in the nutrient removal mechanisms in this study.

### Comparison of bacterial communities

Phyla distributions describing microbial diversity in the BF-A/O-MBR system (sponge biomass and suspended sludge samples) and the C-A/O-MBR system (suspended sludge sample) are summarized in Fig 4(a). A total of 30 phyla were seen from the DNA of the sponge biomass sample of the BF-A/O-MBR system, and 29 phyla were seen from the DNA of the suspended sludge samples of both the BF-A/O-MBR and C-A/O-MBR systems. The term ‘unclassified’ is used to refer to the sequences that could not be classified up to the phylum level. Phyla that were observed at less than 2% average abundance were grouped in ‘Minor phyla’. High proportions of unclassified sequences in a study applying pyrosequencing have been previously reported [17,18]. In this study, depending on the samples, the proportions of the unclassified phyla sequences were 3.06% in the sponge biomass of the BF-A/O-MBR system and, 5.28 and 11.78% in the suspended sludge of the BF-A/O-MBR and C-A/O-MBR systems, respectively.

**Fig 4.**
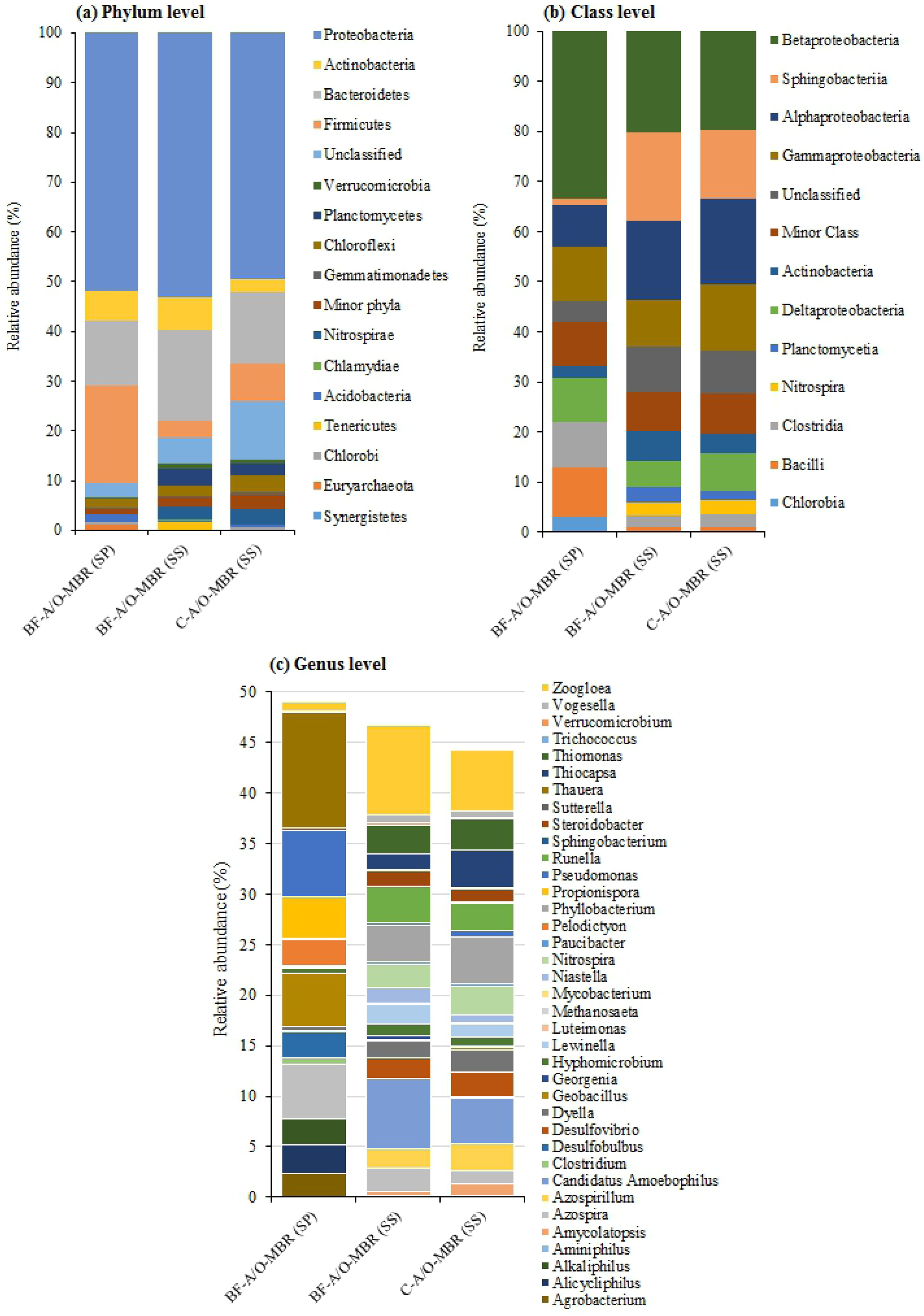
Relative abundance of bacterial phyla level (a), class level (b) and genus level (c) in the sponge biomass sample of BF-A/O-MBR system (BF-A/O-MBR (SP)) and suspended sludge samples of both BF-A/O-MBR and C-A/O-MBR system (BF-A/O-MBR (SS) and C-A/O-MBR (SS), respectively).

According to the pyrosequencing data, about half of the total bacteria in the sponge biomass sample of the BF-A/O-MBR system and the suspended sludge samples of the BF-A/O-MBR and C-A/O-MBR systems belonged to Proteobacteria (51.94, 53.16 and 49.58%, respectively). The results revealed a similar predominant population at the phylum level with other research studies [19,20]. Proteobacteria, a group of Gram-negative bacteria, were found to easily attach themselves to the surface of the media due to the fact that the major components of their outer surface include lipopolysaccharides [21]. Tang et al. [22] mentioned that an adhesive phase was a more suitable environment for the growth of Proteobacteria than that of a suspended phase. Therefore, the sponge could promote the growth of these phylum bacteria in the reactor, which was the main reason why Proteobacteria overwhelmed a larger proportion in both the sponge biomass and the suspended sludge of the BF-A/O-MBR system. The levels of abundance of Bacteroidetes found in the sponge biomass and suspended sludge samples of the BF-A/O-MBR system were 13.22 and 18.06%, respectively, while 14.16% was found in the suspended sludge sample of the C-A/O-MBR system. Bacteroidetes play crucial roles in degrading organic matters and can potentially release proteinaceous EPS [23]. Thus, a higher abundance of Bacteroidetes in the BF-A/O-MBR system is expected to cause the accumulation of biopolymers in the mixed liquor. This finding is consistent with the higher levels of total EPS concentration found in the BF-A/O-MBR system (Table 1).

A very high abundance of Firmicutes was found in the sponge biomass sample (19.35%), meanwhile only 3.43% was observed in the suspended sludge sample of the BF-A/O-MBR system. In comparison with the C-A/O-MBR system, the sponge biomass appeared to trigger a decrease in the relative proportion of Firmicutes in the suspended sludge of the BF-A/O-MBR system, since the higher abundance of this phylum was observed in the suspended sludge sample of the C-A/O-MBR system (7.72%). It was found that Firmicutes were the main factors that influenced membrane fouling in the MBR system [24]. Moreover, the enrichment of filamentous Chloroflexi in the sponge biomass and suspended sludge samples of the BF-A/O-MBR system (1.87 and 2.29%, respectively) were less than in the suspended sludge sample of the C-A/O-MBR system (3.48%). A dominance of filamentous bacteria can promote filamentous bulking in the suspended sludge. Filamentous bulking can also significantly increase the production of SMPs, which in turn substantially increases the fouling of membranes [25]. When considered with the TMP profiles (Supplementary Fig 1S, supporting information), it was found that the fouling rate in the BF-A/O-MBR system (2.60 kPa/day) was almost two times lower than that of the C-A/O-MBR system (4.24 kPa/day). The findings clearly indicate that the sponge addition in the anoxic tank seems to prolong membrane cycle by reducing Firmicutes and filamentous Chloroflexi bacteria in the suspended sludge. In this regard, Firmicutes and filamentous Chloroflexi were more attracted to the sponge surface than the membrane surface.

It was found that the sponge biomass and suspended sludge samples of the BF-A/O-MBR system contained a population of Actinobacteria which accounted for 5.89% and 6.60%, respectively, while in the suspended sludge sample of the C-A/O-MBR system, the figure was 2.62%. Beer et al. [26] has demonstrated that Actinobacteria are important in the biological phosphorus removal processes, due to an ability of anaerobic substrate assimilation and the subsequent aerobic phosphate assimilation and poly-P storage. These results suggest that the BF-A/O-MBR system could achieve higher TP removal efficiency levels when compared with the C-A/O-MBR system by promoting the growth of Actinobacteria in the system. The anaerobic area was created inside the sponge due to the decreasing trend of DO concentration from the surface to the inside of the sponge. Therefore, Actinobacteria may grow in the deep zone of the sponge biomass. When the sponge biomass became thick and was then sloughed off, it could also lead to an increased population of Actinobacteria in the suspended sludge. Nitrospirae was also detected in the suspended sludge samples at a proportion of 2.50 and 2.97% in the BF-A/O-MBR and C-A/O-MBR systems, respectively. However, only 0.06% was observed in the sponge biomass sample. The Nitrospirae bacterium, rather than Nitrobacter, was able to oxidize NO_2_-N into NO_3_-N as has been described in a recent research study [27]. The stable proportion of this phyla might explain why higher and more stable nitrification performance occurred in both systems. The results also indicate that nitrification tended to happen at the outer side rather than the inner side of the sponge biomass.

At the class level, Betaproteobacteria (33.45, 20.08 and 19.58%), Alphaproteobacteria (8.31, 15.86 and 17.10%), Gammaproteobacteria (11.11, 9.36 and 13.28%) and Deltaproteobacteria (8.65, 5.28 and 7.64%) were the most abundance classes in all samples (sponge biomass sample of the BF-A/O-MBR system and suspended sludge samples of the BF-A/O-MBR and C-A/O-MBR systems, respectively) (Fig 4(b)). Betaproteobacteria, the most abundant class, is largely responsible for organic and nutrient removal [28]. Betaproteobacteria, Alphaproteobacteria, Gammaproteobacteria, and Deltaproteobacteria are also well-known for being present in butyrate, glucose, propionate, and acetate-utilizing microbial populations [29]. Alphaproteobacteria, which were a dominant class in some samples of biofilm, are responsible for the biodegradation of some micro-pollutants, i.e., nitrogen and COD removal [30]. The high nitrogen removal in the BF-A/O-MBR and C-A/O-MBR systems (>80%) might be a response of this bacteria group becoming enriched in both systems. Gammaproteobacteria have been shown to favorably adhere to membrane surfaces when compared to those of other microorganisms [31]. A higher relative abundance of Gammaproteobacteria in the suspended sludge sample in the C-A/O-MBR system might be partly responsible for severe membrane bio fouling in the system when compared to the BF-A/O-MBR system.

The relative abundance of Nitrospira and Clostridia was similar in the suspended sludge samples of the BF-A/O-MBR system (2.50 and 2.37%) and C-A/O-MBR system (2.97 and 2.43%), respectively. However, while Nitrospira was very low (0.05%), Clostridia revealed higher abundance (9.13%) in the sponge biomass sample of the BF-A/O-MBR system. Nitrospira plays a pivotal role in nitrification as an aerobic chemolithoautotrophic nitrite-oxidizing bacterium. Nitrospira belongs to the nitrite oxidizing bacteria category, and Nitrospira-like bacteria have been reported as dominant nitrite oxidizers in biofilms that were collected from wastewater treatment plants [32]. Nitrospira and similar bacteria are slow-growing organisms which may be favored for long SRT in this study. Therefore, the suspended sludge might contribute to establishing favorable growth conditions for Nitrospira-like bacteria rather that the sponge biomass, thereby partly contributing to ammonia oxidation in the MBR. As for Clostridia, they are well known to be electrochemically active and capable of fixing N_2_ [33].

A close investigation at the genus level (Fig 4(b)) showed that the genus *Nitrospira,* which is defined as NOB [27], was detected in the suspended sludge samples at proportions of 2.40 and 2.80% in the BF-A/O-MBR and C-A/O-MBR systems, respectively. However, only 0.02% was found in the sponge biomass sample of the BF-A/O-MBR system. Obviously, nitrifiers are likely to prefer a suspended stage rather than an adhesive stage. However, in both systems, the nitrification process was nearly complete and most of the influent NH_4_^+^-N entering the aerobic MBR tank were oxidized entirely into NO_3_ ^-^-N regardless of whether or not a sponge was added. Moreover, the extremely long SRT applied in this study was advantageous for slow-growing bacteria such as nitrifying bacteria [34]. This finding revealed that most of the nitrification process tended to occur outside the sponge biomass and that the existence of a compacted sponge was not involved in the ammonia removing process that occurred in this study. Additionally, pyrosequencing analysis showed that only a small amount of nitrifiers (Fig 5) were detected in the reactor when compared with the qPCR results on the same day, as has been described above (Fig 2). The results suggested that real-time PCR using a functional gene is an effective tool for quantitative targeting of the bacteria group rather than pyrosequencing.

**Fig 5.**
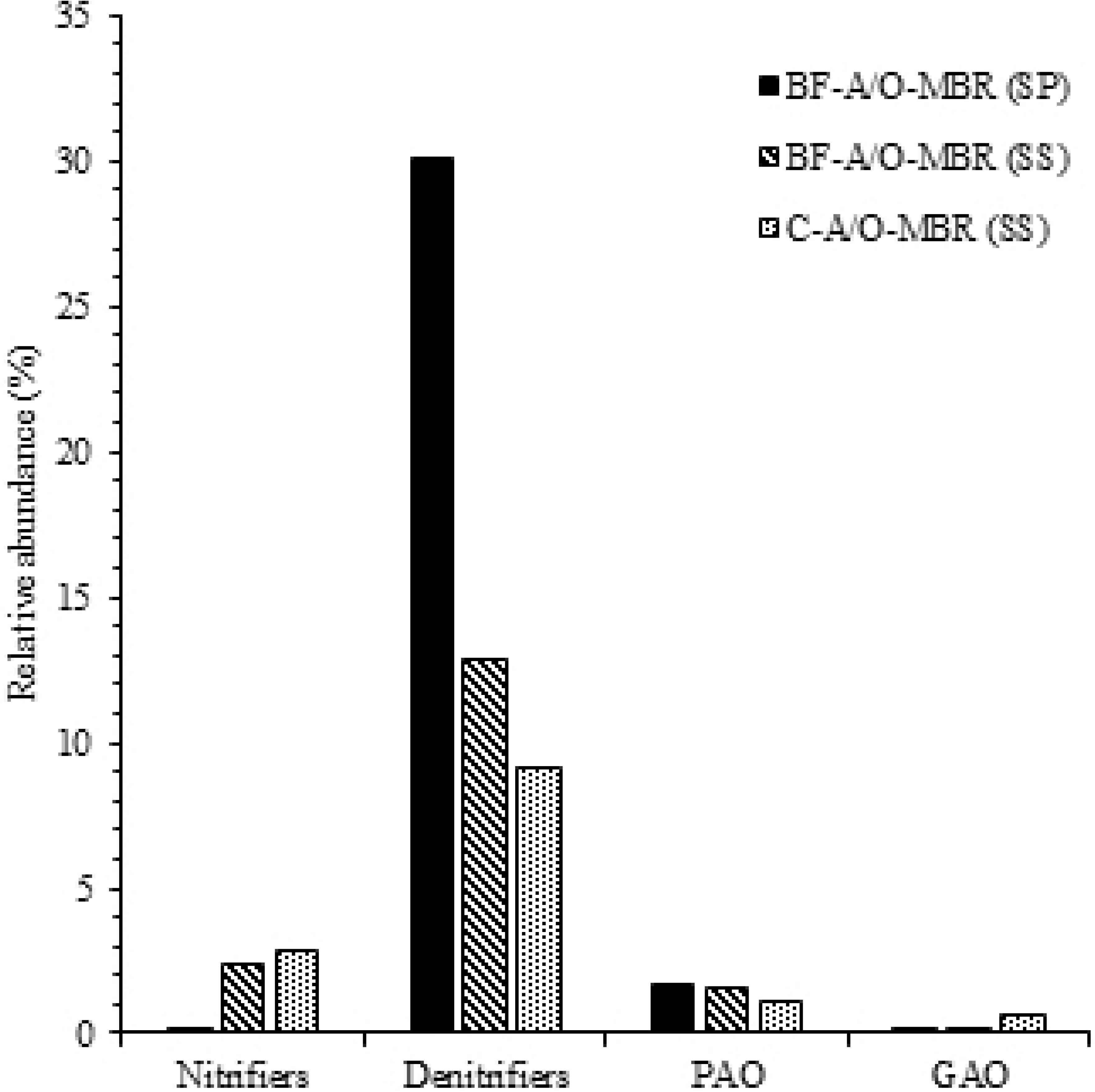
**Relative abundance of potential functional bacterial groups including:** nitrifiers (*Nitrospira*), denitrifiers (*Azospira, Geobacillus, Hyphomicrobium, Pseudomonas, Thauera* and *Zoogloea*), polyphosphate accumulating organisms (PAO: *Dechloromonas*) and glycogen accumulating organisms (GAO: *Sphingomonas* and *Amaricoccus*).

As for denitrifying the bacteria population, six genera were detected in all samples including *Azospira* [35], *Geobacillus* [36], *Hyphomicrobium* [37], *Pseudomonas* [38], *Thauera* [39] and *Zoogloea* [34]. As is shown in Fig 5, the highest level of abundance of denitrifying bacteria was found in the sponge biomass sample of the BF-A/O-MBR system, accounting for 30.10%. However, they accounted for a lower proportion in the suspended sludge sample of the BF-A/O-MBR and C-A/O-MBR systems (12.08 and 9.09%, respectively). The highest genus in the sponge biomass sample was *Thauera* (11.40%) followed by *Pseudomonas* (6.59%), *Azospira* (5.43%) and *Geobacillus* (5.33%), while the highest genus in the suspended sludge samples of the BF-A/O-MBR and C-A/O-MBR systems was *Zoogloea* (8.86 and 6.02%, respectively) followed by *Azospira* (2.32 and 1.20%, respectively) and *Hyphomicrobium* (1.18 and 0.94%, respectively). The deficiency of denitrifying bacteria in the C-A/O-MBR system can be a reflex of the significantly lower value of TN removal efficiency (α = 0.05). A significantly higher abundance of denitrifying bacteria in the sponge biomass sample revealed that the presence of a compacted sponge in the BF-A/O-MBR system could improve TN removal by promoting the growth of some of denitrifying bacteria groups in the reactor. However, the pyrosequencing results of the denitrifying bacteria (Fig 5) are in contrast to the qPCR results (Fig 2). As mentioned above, qPCR results that had been achieved with a functional gene showed that the abundance of denitrifying bacteria in the sponge biomass sample of the BF-A/O-MBR system was lower than in the suspended sludge samples of both the BF-A/O-MBR and C-A/O-MBR systems. The results indicated that some of the denitrifying bacteria detected by pyrosequencing in the sponge biomass sample might not have any ability to denitrify nitrates in this study.

*Dechloromonas,* which have the ability to store polyphosphates (PAO) in full-scale wastewater treatment plants [40], were found to be predominant in both the sponge biomass and suspended sludge samples of the BF-A/O-MBR system (1.71 and 1.51%, respectively) rather than the suspended sludge samples of the C-A/O-MBR system (1.10%). *Sphingomonas* and *Amaricoccus* related glycogen accumulating organisms (GAOs) [41,42] were also detected in all samples. The abundance of these two genera was found in the suspended sludge sample of the C-A/O-MBR system (0.44 and 0.25%, respectively). However, lower than 0.1% was observed in both the sponge biomass and the suspended sludge sample of the BF-A/O-MBR system. As has been illustrated in Fig 5, PAOs and GAOs seemed to compete with each other. Numerous studies have shown that GAOs and PAOs compete for the same substrate under anaerobic conditions, but no accumulation of polyphosphates occurred under aerobic conditions [43]. These results indicate that the absence of a sponge biomass inhibited the growth of PAOs and increased the abundance of GAOs resulting in the deterioration of the phosphorus removal performance. Moreover, this finding also confirmed the hypothesis that the sponge media could provide an anaerobic environment under anoxic conditions for PAOs to consume a carbon source and release phosphates at the same time, resulting in a high degree of phosphorus removal that was achieved even without the anaerobic compartment.

To obtain a higher resolution of community composition, a heatmap was utilized to illustrate the relative abundance of the 37 OTUs found in the three samples presented in Fig 6. Three major groups were identified and were separated by only six units in the bacterial community pattern cluster analysis, indicating clear distinctions of microbial community structure due to the different stages that exist (adhesive and suspended stage) and with and without the addition of a compacted sponge. As is shown in Fig 6, clusters 1 and 2, which are comprised of *Azospirillum, Desulfovibrio, Nitrospira, Phyllobacterium, Runella, Thiocapsa, Thiomonas, Dyella, Hyphomicrobium, Lewinella, Niastella, Steroidobacter* and *Vogesella,* were dominant in the suspended sludge sample (BF-A/O-MBR (SS) and C-A/O-MBR (SS)) of both systems. Most of them belong to phyla Proteobacteria and Bacteroidetes which are commonly found in activated sludge and biofilm. The heatmap clearly shows that the existence of compacted sponge in the BF-A/O-MBR system could promote the growth of *Zoogloea, Candidatus Amoebophilus* and *Azospira* (cluster 5) in the suspended sludge. A significant difference in the microbial community was observed in the sponge biomass sample of the BF-A/O-MBR system (BF-A/O-MBR (SP)). The levels of abundance of clusters 1 and 2 decreased in the suspended sludge samples, while levels in clusters 4 and 6 were more dominant in the sponge biomass sample of the BF-A/O-MBR system. *Thauera, Propionispora, Geobacillus, Pseudomonas,* and *Azospira* were found to be distinctly present in the sponge biomass sample. Due to the denitrification capacities of *Zoogloea, Azospira, Thauera, Geobacillus* and *Pseudomonas,* which are dominant in the BF-A/O-MBR system, indicated that a significantly higher level of TN removal that was found in this system was encouraged by the presence of the compacted sponge inside the anoxic tank.

**Fig 6.**
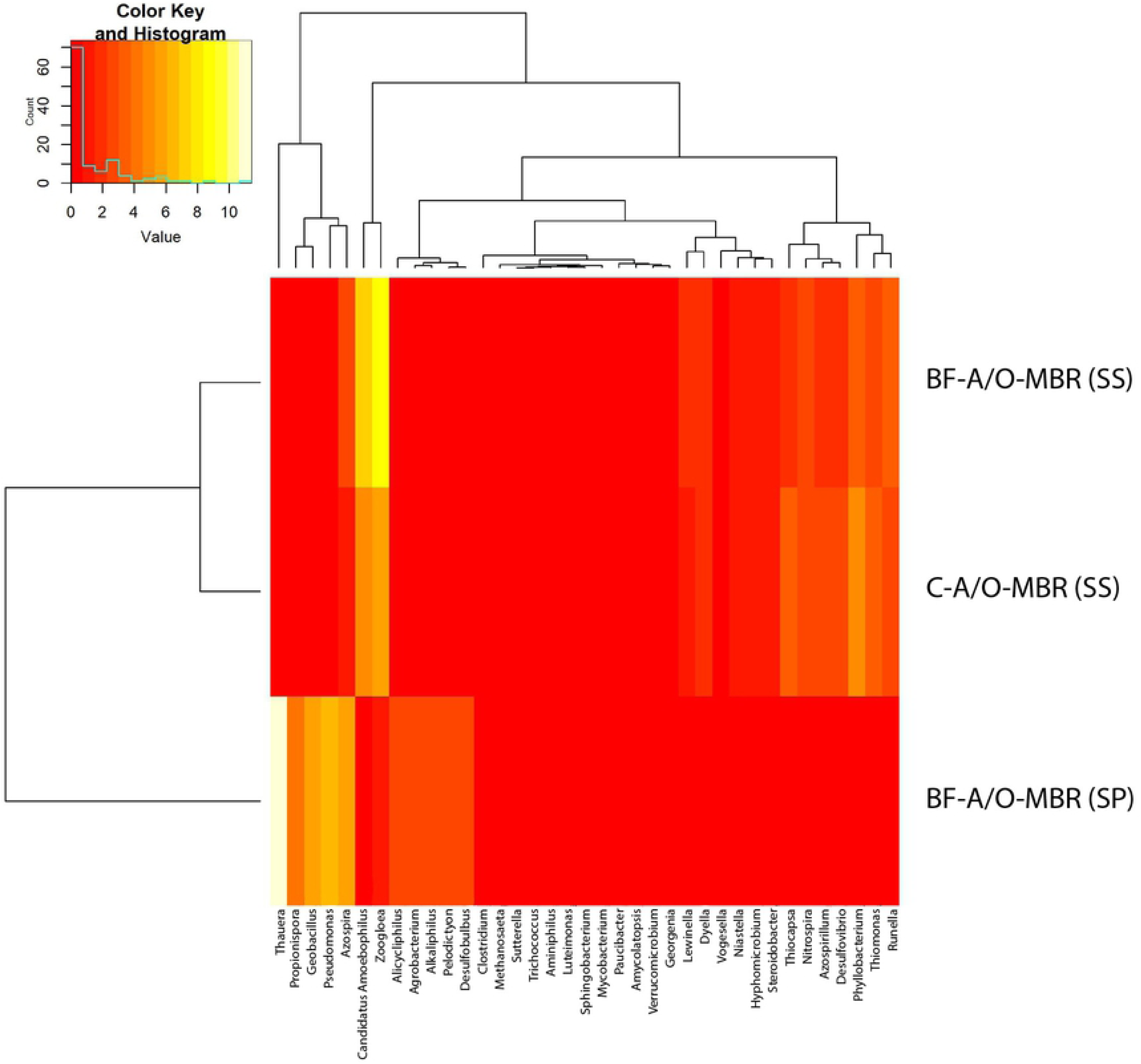
Heatmap of cluster analysis of the most abundant genera (>2% of the total OTU reads in each group) for sponge biomass sample of BF-A/O-MBR system (BF-A/O-MBR (SP)) and suspended sludge samples of both BF-A/O-MBR and C-A/O-MBR system (BF-A/O-MBR (SS) and C-A/O-MBR (SS), respectively) (b).

### Correlation between nutrient removal and bacterial community

To investigate the ecological correlation between the bacterial community composition (at the genus level), and nitrogen removal, Pearson’s correlation was conducted using TKN, organic-nitrogen (Org-N), ammonia-nitrogen (NH_4_), nitrite-nitrogen + nitrate-nitrogen (NOx) and total nitrogen (TN) concentrations in different zones (anoxic tank and MBR tank) as key variables (Fig 7). A heatmap of Pearson’s correlation showed that the genus *Amycolatopsis* (r_p_ = −0.94), *Azospirillum* (r_p_ = −0.96), *Candidatus Amoebophilus* (r_p_ = −0.94), *Desulfovibrio* (r_p_ = −0.99), *Dyella* (r_p_ = −0.97), *Georgenia* (r_p_ = −0.84), *Hyphomicrobium* (r_p_ = −0.94), *Lewinella* (r_p_ = −0.95), *Luteimonas* (r_p_ = −0.93), *Nitrospira* (r_p_ = −0.99), *Paucibacter* (r_p_ = −0.97), *Phyllobacterium* (r_p_ = − 0.98), *Runella* (r_p_ = −0.97), *Sphingobacterium* (r_p_ = −1.00), *Steroidobacter* (r_p_ = −0.99), *Thiocapsa* (r_p_ = −0.81), *Thiomonas* (r_p_ = −1.00), *Trichococcus* (r_p_ = −0.90), *Vogesella* (r_p_ = −0.99) and *Zoogloea* (r_p_= −0.94) contributed to the reduction of total nitrogen in the system due to a strong negative correlation with total nitrogen concentration. The genus *Nitrospira* was closely related to total nitrogen removal in the system owing to nitrification, while *Hyphomicrobium* and *Zoogloea* were assigned to total nitrogen removal in the system owing to denitrification. Therefore, it could be concluded that among the six denitrifying bacteria (i.e., *Azospira, Geobacillus, Hyphomicrobium, Pseudomonas, Thauera* and *Zoogloea*) detected in this study, *Hyphomicrobium* and *Zoogloea* were the most prevalent genera and that this outcome was related to the denitrification process. *Hyphomicrobium* and *Zoogloea* were most abundant in the suspended sludge samples of the BF-A/O-MBR system, followed by the suspended sludge sample of the C-A/O-MBR system. However, they were found to be lacking in the sponge biomass sample of the BF-A/O-MBR system. This finding clearly revealed that denitrifying bacteria, which have an important ability in the denitrifying process, preferred a suspended stage rather than an adhesive stage for growth. However, the presence of a compacted sponge in the BF-A/O-MBR system seemed to promote the growth of denitrifying bacteria when compared with the conventional C-A/O-MBR system. As for other genera, the findings are still unclear due to a lack of information regarding these genera and their relationship to nitrogen removal. Due to the fact that the process of nitrogen removal could occur by simultaneous biomass assimilation and dissimilation (nitrification-denitrification process), a possible reason for this is that these bacteria genera may contribute to total nitrogen removal by cell assimilation.

**Fig 7.**
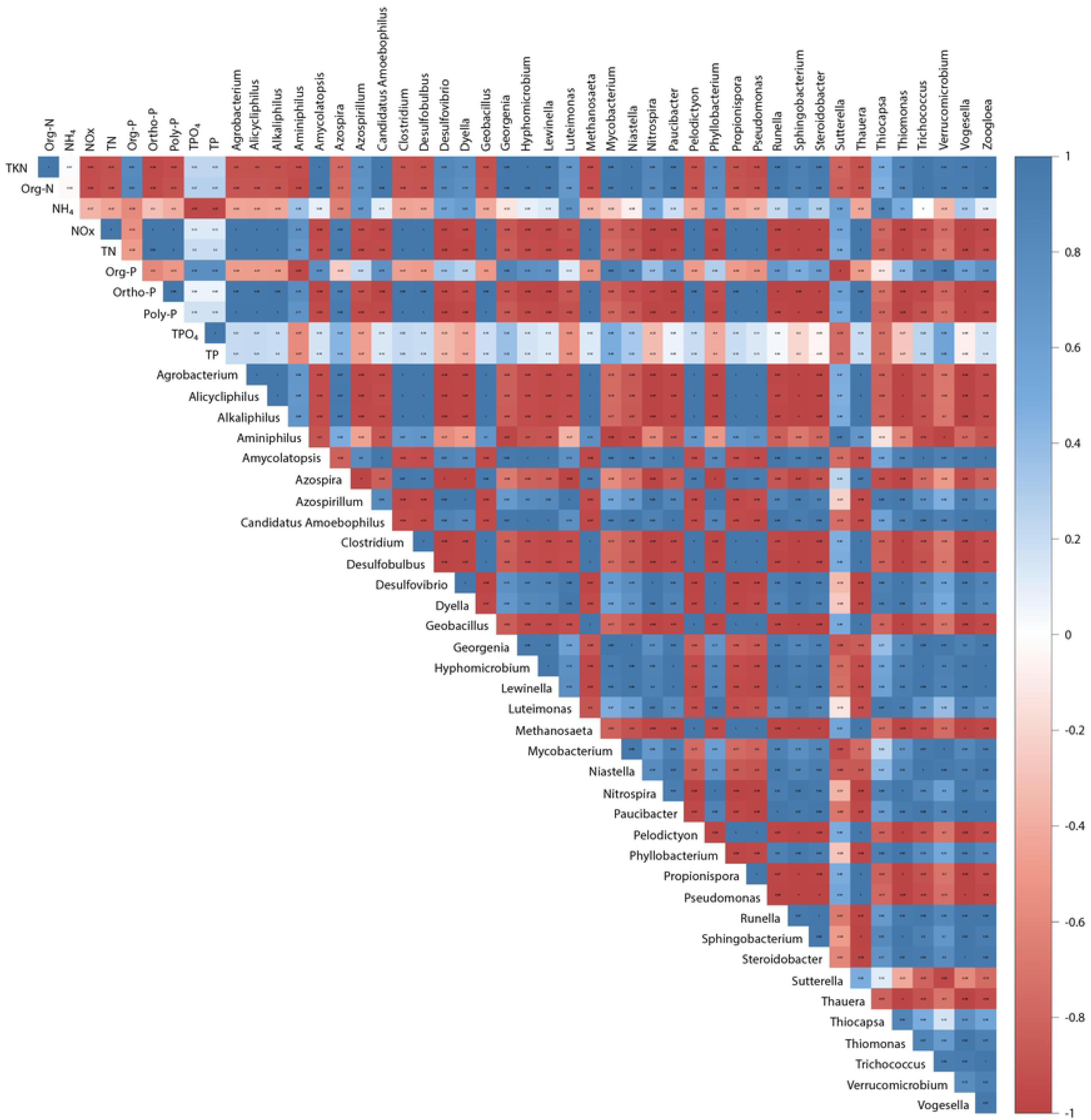
Heatmap of Pearson’s correlation between the most abundant bacteria (>2% of the total OTU reads in each group) and nutrients concentration in the system.

A heatmap of Pearson’s correlations between bacterial community composition (at the genus level) and phosphate forms is also shown in Fig 7. The results show that the correlation coefficient values (r_p_) between *Azospira* and polyphosphates, and total phosphates were found to be 0.96 and 0.98, respectively, which indicated that the impact of genus *Azospira* has a strong positive correlation with poly-phosphate and total phosphate levels. Although several reports have identified the genus *Azospira* as a denitrifying bacterium in wastewater treatment plants [35,44], some species belonging to the genus of *Azospira* were recently identified and are also considered to be potential PAOs [45]. Since both systems operate under complete SRT, the microorganisms that have the ability to accumulate phosphates did not withdraw from the reactor and most of them were recirculated within the reactor until the end of the experiment. The abundance of PAOs coupled with an increase of phosphate concentration in the reactor (positive correlation) may be involved with the phosphorus removal mechanisms in this study. Therefore, *Azospira,* which revealed higher levels of abundance in the sponge biomass (5.43%) and suspended sludge (2.32%) of the BF-A/O-MBR system when compared with the C-A/O-MBR system (1.20%), seemed to play a role in phosphorus removal in this study. On the contrary, *Thiocapsa* was found to have a significantly negative correlation with orthophosphate (r_p_ = −0.73), poly-phosphate (r_p_ = −0.79) and total phosphate (r_p_ = −0.73) concentrations. This indicated that as the abundance of the genus *Thiocapsa* increased, lower levels of orthophosphate, poly-phosphate and total phosphate concentrations accumulated in the reactor. However, the investigation is still unclear and there is a lack of information that can indicate what the main mechanism of phosphorus in the BF-A/O-MBR system is, when operated with complete SRT. Further studies need to be performed to clearly understand the mechanisms of this phenomenon.

## Conclusions

Two A/O-MBR systems (BF-A/O-MBR and C-A/O-MBR) were operated in parallel to compare system performance and microbial community composition. High average removal of COD, NH_4_^+^-N and TN was achieved in both systems. However, TP removal efficiency was remarkably higher in BF-A/O-MBR when compared with the C-A/O-MBR. TP mass balance suggested that under complete SRT, sponges play key roles in phosphorus release and accumulation. Results of pyrosequencing clearly reveal that a compacted sponge in BF-A/O-MBR could promote the growth of bacteria involved in nutrient removal and reduce the growth of filamentous and bacteria related to membrane fouling in the suspended sludge.

## Acknowledgments

The authors gratefully acknowledge the financial support provided by Kurita-AIT Research Grant. The research work was done during the study period of Ph.D. degree at Chiang Mai University. The authors would also like to extend their gratitude to the Graduate School, Chiang Mai University, for the financial support. This research work was partially supported by Chiang Mai University

## Supporting information

**1S Table. Oligonucleotide primers used for amplification.**

**2S Table. Reagents mixture composition for qPCR Method: SYBR Green 1.**

**3S Table. Phosphorus mass balance analysis in BF-A/O-MBR and C-A/O-MBR system.**

**1S Fig. Trans-membrane pressure (TMP) profiles of the A/O MBR systems.**

